# Towards EEG-to-Text: Handwritten Character Classification via Continuous Kinematic Decoding

**DOI:** 10.1101/2024.04.26.591369

**Authors:** Markus R. Crell, Gernot R. Müller-Putz

## Abstract

The classification of handwritten letters from invasive neural signals has lately been subject of research to restore communication abilities in people with limited movement capacities. This study explores the classification of ten letters (*a,d,e,f,j,n,o,s,t,v*) from non-invasive neural signals of 20 participants using two methods: the direct classification from low-frequency and broadband electroencephalogram (EEG) and a two-step approach comprising the continuous decoding of hand kinematics and the application of those in subsequent classification. When using low-frequency EEG, results show moderate accuracies of 23.1 % for ten letters and 39.0 % for a subset of five letters with highest discriminability of the trajectories. The two-step approach yielded significantly higher performances of 26.2 % for ten letters and 46.7 % for the subset of five letters. Hand kinematics could be reconstructed with a correlation of 0.10 to 0.57 (average chance level: 0.04) between the decoded and original kinematic. The study shows the general feasibility of extracting handwritten letters from non-invasively recorded neural signals and indicates that the proposed two-step approach can improve performances. As an exploratory investigation of the neural mechanisms of handwriting in EEG, results suggest movement speed as the most informative kinematic for the decoding of short hand movements.

## Introduction

The restoration of motor capabilities for patients with impaired neuromuscular function has recently received increasing attention in scientific research resulting in significant advances in this area. Individuals with spinal-cord injuries are inhibited primarily in their execution of daily activities and require, among other things, the restoration of actual movement execution. Patients with locked-in syndrome or similar medical conditions, however, additionally experience severe impairments in their communication abilities and require foremost reliable methods for the interaction with relatives and caretakers (***Vansteensel et al., 2023***). Over the last years, the restoration of comunication channels for these patients has seen extensive progress driven by advances in the field of brain-computer interfaces (BCIs). Early BCIs for communication adopted matrix-based spellers to select characters based on P300 paradigms of sequential flashing of rows/columns (***Polich, 2007***) or sensorimotor-based spellers decoding discrete movement attempts for the selection of characters (***Neuper et al., 2006***). The expandability of sensorimotor-based BCIs to multiple classes of movement like wrist flexion/extension or hand opening/closing (***Ofner et al., 2017a***) or the detection of movement direction in center-out tasks (***Kobler et al., 2020a; Shiman et al., 2017***) further enhanced the capabilities for character selection or enabled the step-wise control of end-effectors (***Schwarz et al., 2020; Jeong et al., 2018***). A wide range of motor-based BCIs is based on movement-related cortical potentials (MRCPs). These neural correlates of movement are phase- and time-locked to the movement onset (***Shibasaki and Hallett, 2006***) and encode information about the type of movement (***Ofner et al., 2017b; Iturrate et al., 2018***) and parameters like force (***Jochumsen et al., 2013***) or speed (***Gu et al., 2009; Xu et al., 2021***). MRCPs occur in low-frequency time-domain EEG and exhibit a negative deflection starting up to 2 s before the movement onset. They reach the maximum negativity at the start of the movement and show a positive deflection around 300 ms after the movement (***Shibasaki and Hallett, 2006***). The maximum negative deflection is reached in central motor areas contralateral to the movement. Several studies have shown that the classification of discrete movements from MRCPs is possible in both healthy (***Pereira et al., 2021; Xu et al., 2021***) and disabled people (***Ofner et al., 2019***).

Recently, attempts to shift from discrete classification of single movements to continuous control of end-effectors such as computer cursors in non-invasive BCIs have substantially progressed. ***Mondini et al. (2020***) implemented trajectory decoding of hand movements for the online control of a robotic arm and, following this, were able to improve the decoding accuracy by incorporating non-directional kinematics such as distance and speed (***Kobler et al., 2020c***). Their approach utilized partial least squares (PLS) regression for the inference of movement kinematics from electroencephalography (EEG) signals and an Unscented Kalman Filter (UKF) to combine the decoded kinematics into one trajectory. ***Kobler et al. (2020c***) showed that non-directional kinematics are encoded differently in neural data and can offer additional information about the movement which can be utilized to improve the decoded motion. ***Pulferer et al. (2022***) validated this approach in a study including a spinal-cord injured participant. The inclusion of error potentials into the cursor-control system further improved the performance of the system (***Mondini et al., 2024***), showing the general feasibility of trajectory decoding from non-invasive neural signals. Other methods including the combination of convolutional neural networks (CNNs) and long short-term memory (LSTM) architectures have also been utilized for the reconstruction of continuous hand trajectories where they have been shown to yield high performances for the decoding of unimanual three-dimensional (***Jeong et al., 2020***) and bimanual trajectories (***Chen et al., 2022***). A direct comparison (***Borra et al., 2023***) between multiple models for two-dimensional unimanual trajectory decoding including CNNs, a CNN+LSTM combination and traditional machine learning approaches showed that CNNs outperformed most other methods and performed similar to the PLS+UKF approach implemented by ***Kobler et al. (2020c***).

Significant progress has further been accomplished in the area of re-enabling communication with intracortical and intracranial BCIs which utilize invasive recordings of neural signals from electrode grids or microelectrode arrays. Movement or communication restoration through single- (***J. et al., 2016***) and multi-class detection (***Schwemmer et al., 2018; Bouton et al., 2016***), inference of words from attempted speech (***Metzger et al., 2023; Willett et al., 2023; Moses et al., 2021***) and the restoration of uni- and bimanual effector control (***Deo et al., 2024; Simeral et al., 2020; Nuyujukian et al., 2018***) show the extensive advances that have been reached in this field. The high performance of communication that is enabled through such methods is evident in a recent communication system proposed by ***Willett et al. (2021***) which currently constitutes one of the fastest and most accurate communication channels for locked-in patients. Their approach utilizes the classification of handwritten characters from attempted movement and achieves a speed of 90 characters per minute with an accuracy of 94.1 %. Integrating a language model for the correction of spelled words, the accuracy could be increased to 99 %. While the authors did not utilize the decoding of motion trajectories for the classification of letters, they showed the possibility of reconstructing the written letters from neural signals and provided evidence for a tuning of neural data to the speed of the writing motion.

Although this decoding of handwritten letters from intracortical neural signals has been shown to yield exceptional results enabling high-speed communication, similar approaches have received limited interest for the usage with non-invasive BCIs. ***Pei and Ouyang (2021***) decoded handwritten letters non-invasively from five participants during repetitive writing of a fixed phrase containing nine different symbols. While they achieved a high accuracy of 76.8–97.0 %, the authors did not correct for eye artifacts prior to the classification, thereby possibly biasing their results.

In the current study, we aimed to explore the neural mechanisms and classification of handwritten letters from non-invasive neural signals. Based on previous research on invasive classification of handwritten letters, we hypothesized that neural differences between letters would similarly occur in non-invasive signals and that letters would be successfully discriminated based on EEG recordings. Due to the commonly observed reduction in accuracy when discriminating multiple movement classes from EEG, we aimed to compare two different approaches for the classification: (a) a direct approach classifying the letters from low-frequency and broadband EEG data and (b) a two-step approach of decoding continuous hand kinematics from neural data and classifying the letters based on the reconstructed trajectories. We hypothesized that, based on promising advances in the non-invasive decoding of kinematics in other studies and previously reported tuning of neural data to kinematics during handwriting, the proposed two-step approach would lead to increases in the classification performance.

## Materials and Methods

### Participants and Setup

Twenty-two healthy, right-handed subjects (eleven male, eleven female) with a mean age of (mean ± standard deviation) 27.5 ± 3.92 years and normal or corrected-to-normal vision participated in the study. Participants gave written informed consent before partaking and received monetary compensation for their time. Two subjects had to be excluded from the study, one due to technical problems during the recording and the other due to early withdrawal from the experiment. All participants were informed of the study procedure and potential risks and subsequently equipped with the measurement setup consisting of 60 EEG electrodes and four electrooculography (EOG) electrodes positioned about 2 cm away from the lateral canthi of the eyes and above and below the left eye. The EEG channels were positioned according to a standard 10-10 montage with the exclusion of channels F9, F10, T9, T10, TP9, TP10, P9 and P10 and additional channels at PPO1h and PPO2h. EEG and EOG data was recorded with biosignal amplifiers (BrainAmp, Brain Products GmbH, Germany) at a rate of 500 Hz. The measurement setup further consisted of a motion capture system to record the position of the right index finger. The system used a camera mounted above the hand and an optical marker which was attached to the index finger to record 2D positions at a rate of 30 Hz. Participants executed all movements inside a specifically designed box which incorporated the motion capture system and a haptic border to ensure that participants did not move their fingers outside the monitored area. During the movements, the hand of the participant was positioned inside the box and the view of the hand was obstructed with a black cloth to ensure that no eye movements related to the motion of the hand could occur. The complete setup together with the motion capture box and the screen on which the paradigm was displayed is shown in Figure 1a.

**Figure 1.**
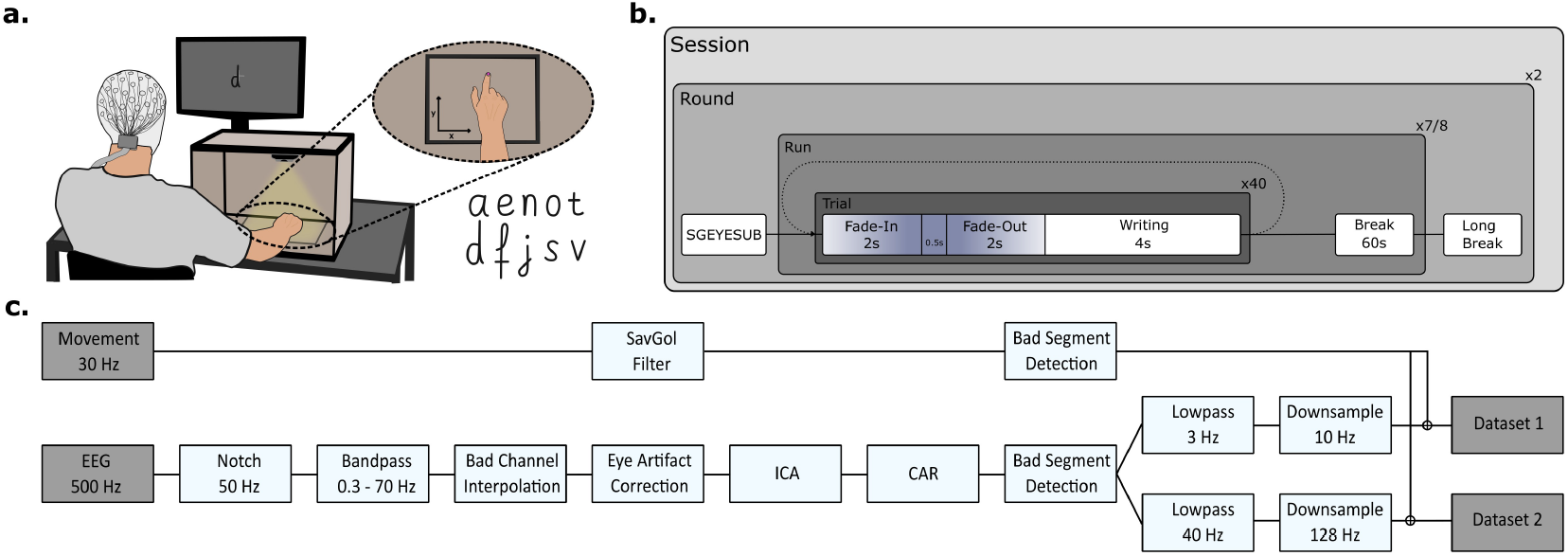
(a) The experimental setup consisting of a display and a motion capture system positioned inside a box for obstruction of vision during hand movements. A haptic border inside the box ensured that the participants did not move their finger outside of the monitored area. Ten different letters as given on the bottom right were shown on the display during the paradigm. (b) Timeline of the handwriting paradigm as executed by the participants during the measurement. In total, participants wrote 600 letters over two rounds with a duration of approximately 45 min each. The fist round consisted of seven runs while the second round consisted of eight runs. The duration of the total experiment was approximately 150 min. (c) Preprocessing pipeline used for the processing of the recorded neural and kinematic data.

### Experimental Paradigm

EEG data was collected during a paradigm in which subjects were tasked with the writing of different letters using the index finger of their right hand. A complete session consisted of an instruction phase, a training phase and two measurement rounds with a break of 5–10 min between them. Subjects were seated comfortably in a chair in front of a display showing the instructions and the paradigm. Prior to each round, specified eye movements were recorded to enable eye artifact correction of the recorded data. A measurement round consisted of seven (first round) or eight (second round) writing runs separated by one minute breaks during which the participant was allowed to move and relax. Each run contained 40 trials with a duration of 8.5 s each. At the beginning of a trial, the letter that participants were supposed to draw faded onto the center of the screen over a duration of 2 s (*fade-in* phase). It remained at full opacity for 0.5 s and faded out again over a duration of 2 s (*fade-out* phase). Participants were instructed to start their movement as soon as the letter was faded out entirely. A *writing* phase of 4 s during which the letter was supposed to be written completely followed and the next trial started immediately afterwards. During the *writing* phase, only a fixation cross was presented to the participants to ensure that no eye movements related to the written letter would occur. We specifically designed the study paradigm to minimize eye movements during movement execution based on findings from ***Kim et al. (2015***) on the influence of eye artifacts during trajectory decoding. Participants were instructed to stop their hand at the last position of the letter after they finished writing until the next letter would fade onto the screen. The fading of the letter as the cue for the start of the movement was selected to minimize the influence of visual processing on the neural data as has been shown by ***Ofner et al. (2019***). Except for the first trial in every run, participants were instructed to execute a *home movement* during the *fade-in* phase where they moved their hand back to a comfortable position. This *home movement* was necessary to ensure a neutral starting position and to avoid that participants would draw outside of the boundary of the monitored area. During one run, ten different letters were shown to the participant in random order. This was repeated four times per run, resulting in a total duration of 340 s per run. Over the total of 15 runs, each letter was written 60 times. The displayed and written letters were *a,d,e,f,j,n,o,s,t* and *v* and are shown in Figure 1a. The paradigm structure is displayed in Figure 1b.

To ensure that participants were aware of the intended way of writing the letters and to minimize inter-participant variability in the size and speed of the writing, a training was completed before the rounds. In four steps, the participants would (1) observe the intended trajectory by observing a red dot follow the shape of the letter, (2) follow the red dot and the letter shape by moving their finger and receiving feedback about their finger position on the display, (3) follow the shape of the letter with their finger without any displayed letter but while receiving feedback on the display and finally (4) execute the movement without any feedback on the display. The last step of the training was similar to the actual paradigm and served the purpose of validating the participant’s understanding of the paradigm through the experiment leader.

In combination with the training performed prior to the first round, the execution of each of the two rounds took approximately 45 min with the whole session including preparation and instructions lasting for approximately 150 min.

### Letter Selection

The letters *a,d,e,f,j,n,o,s,t,v* were selected based on two criteria: *a,e,n,o* and *t* were selected as the most frequent lower-case letters in the English alphabet (***Jones and Mewhort, 2004***) based on the final objective of communication through the recognition of handwritten letters. The other letters (*d,f,j,s* and *v*) were selected based on the discriminability of their writing trajectory. To obtain five letters with the most discriminable writing trajectory, the trajectories of 21 letters of the alphabet (without *a,e,n,o,t*) were normalized to the same writing speed and converted to angular trajectories by computing the angle with the x-axis for every point on the trajectory. We calculated the pairwise distance between all letters as the mean difference between the angular trajectories. For letters with different lengths of the trajectory, the maximum distance was used for the remaining time points. Based on this distance matrix, k-means clustering was performed to split the letters into five clusters. From these, the set with the largest overall distance between letters was selected. The difference in angular trajectory was established as a metric for discriminability between letters based on its importance in the perception of shape similarity (***Zhang and Lu, 2002***) and its invariance to the size of a shape.

### Data Preprocessing

The recorded data was preprocessed in Python following specified pipelines for kinematic and neural data. The kinematic trajectory obtained from the motion capture system was smoothed with a Savitzky-Golay filter (first order polynomials and window length of 200 ms). Start (onset) and stop (offset) of the handwriting movements were obtained by calculating the movement speed and detecting the falling and rising edges from the baseline. Neural data was notch filtered at 50 Hz and bandpass filtered between 0.3 and 70 Hz using a zero-phase Butterworth filter of fourth order. Bad channels were visually selected and interpolated from neighboring locations. The sparse generalized eye subspace subtraction (SGEYESUB) algorithm (***Kobler et al., 2020b***) was then used to correct eye artifacts using the collected EOG data. Remaining eye artifacts as well as muscle or sweat artifacts were removed from the data using independent component analysis (ICA) after manual inspection of the components. The data was re-referenced to a common average reference (CAR). Bad segments were identified and marked for rejection upon surpassing a threshold of ±120 µV or when the kurtosis or probability of a segment exceeded seven standard deviations from the mean. Two datasets for usage in the training and validation of different models were created by lowpass filtering the data with a zero-phase Butterworth filter at two different cutoff-frequencies and signals were downsampled accordingly (dataset 1: 0.3–3 Hz, *f*_*s*_ = 10 Hz, dataset 2: 0.3–40 Hz, *f*_*s*_ = 128 Hz). Each set was split into trials of 8.5 s length including 4.5 s before and 4 s after the actual onset of the movement as detected from the motion capture data. Trials which included a rejected segment of neural data, an incomplete movement or in which the wrong letter was written were excluded. Due to these exclusion criteria, an average of 46.5 trials per letter and subject were obtained. The complete preprocessing pipeline for neural and kinematic data is shown in Figure 1c.

### Analysis of Neural Correlates

We investigated whether writing speed or duration of the writing movement influenced the amplitude of the negative deflection of the MRCP associated with the movement onset by correlating the average speed and duration with the amplitude of the subject-specific MRCP in channels C3, C1, Cz, C2 and C4. Subject-specific MRCPs were extracted by averaging the low-frequency EEG data of movement onset centered trials (dataset 1) of the letters per subject to generate an average neural response time-locked to the movement onset. The negative deflection of the subject-specific MRCPs was extracted from the minimum of the amplitude in a range of ±400 ms around the movement onset.

We utilized the average speed and duration of each letter per subject to obtain the slopes of the regression line between negative deflection and speed/duration. An independent t-test was used to determine whether the obtained slope of the regression lines was significantly different from zero. We similarly investigated the correlation between average speed or duration and the rebound rate (***Gu et al., 2009***) defined as the difference in amplitude between the maximum negative deflection ±400 ms around the movement onset and the amplitude 1.0 s later. All p-values were corrected for multiple comparisons using the Bonferroni method.

We further examined neural differences between letters in low-frequency EEG. At every time step in the range of [−0.5; 1.5]s relative to the movement onset, a one-way ANOVA was calculated per channel to check for significant differences between the ten letters. All p-values were corrected for multiple comparisons using the Bonferroni method and, for each pair of *channel* × *time step* with a p-value ≤ 0.01 after correction, a pairwise Tukey’s Honest Significant Difference test was applied to determine which letters showed significant differences in neural data.

### Letter Classification

Letters were classified using a sliding-window approach dependent on the parameters *window length w* and *latency l*. The movement onset centered trials were cut into segments of the specified length at a specific latency which defined the shift of the window relative to the movement onset. For each window length and latency, a set of data segments was created. For each trial, a segment contained the EEG data for [*l* − *w; l*]. Window length and latency were utilized in a defined range of *w* ∈ [0.5, 1.0, 1.5, 2.0, 2.5, 3.0, 3.5]s, *l* ∈ [−0.5; 3.5]s with the latency *l* being advanced with a stride of 0.1 s. For each subject, window length and latency, one model was trained and evaluated such that a classification accuracy was obtained for every model. Models were evaluated using a two-times repeated 5-fold cross-validation strategy. The classification was carried out with two different methods: a shrinkage linear discriminant analysis and a CNN based on the EEGNet architecture.

Linear discriminant analysis (LDA) is a supervised algorithm for finding linear combinations of features to optimally separate the data into a number of predefined classes. Since small datasets, as are common in BCIs, can lead to imprecise estimates of the utilized covariance matrix, we employed shrinkage LDA (sLDA). This method utilizes a regularization parameter γ to shrink the covariance matrix and overcome the imprecise estimation (***Blankertz et al., 2011***). The implementation of sLDA used in the current study employed the quadratic-inverse shrinkage (QIS) estimator (***Ledoit and Wolf, 2019***). sLDA is commonly used for the classification of EEG data in movement-related tasks and has been shown to outperform the classical LDA while also requiring less training data to achieve comparable results (***Lotte et al., 2018***). The classification of movement-related EEG data using sLDA has been shown to produce results with high accuracy when using low-frequency data (***Ofner et al., 2017b***).

Similar to sLDA, CNNs are commonly used in the classification of neural data. ***Lawhern et al. (2018***) proposed a Convolutional Neural Network adapted to classify EEG data for different tasks, the EEGNet, which has been used extensively in the classification of EEG signals. The basis of this model is the automatic feature-selection which finds appropriate spatial filters for the extracted frequency bands directly from training data and thus eliminates the need to manually tailor features specifically to the current task. The model architecture is given in Figure 2b. In this study, we employed standard parameters for the classification of ten different classes. A dropout rate of 0.6 was used to avoid overfitting and the learning rate was adapted with exponential decay throughout the training.

**Figure 2.**
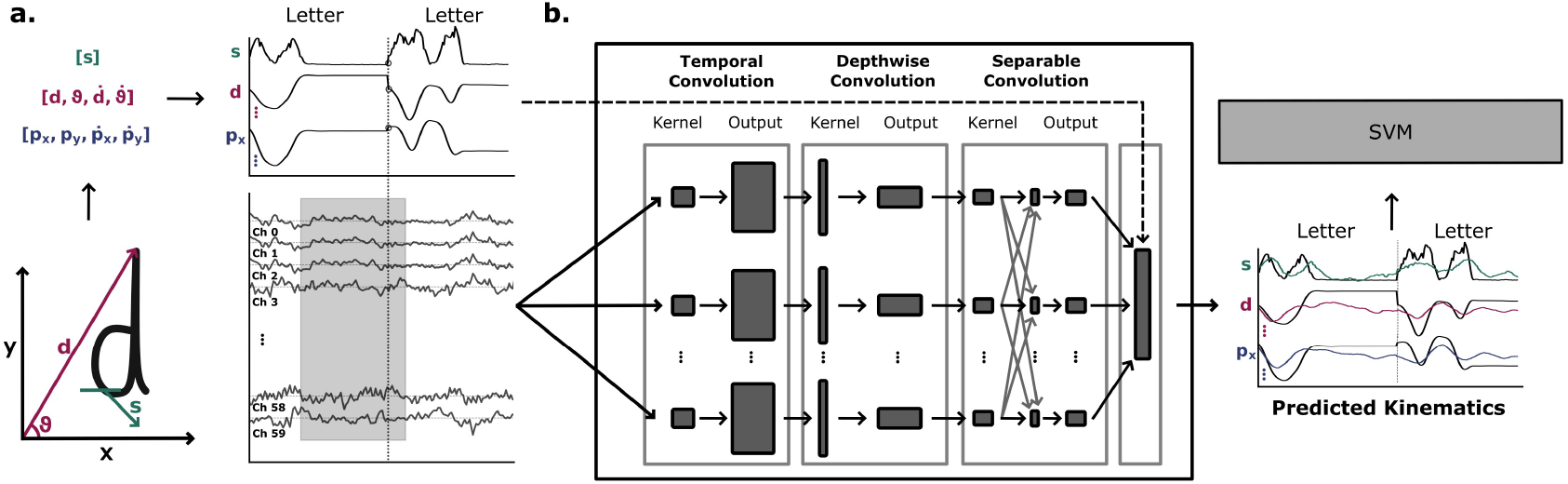
(a) Schematic of the input data for decoding of hand kinematics from EEG. The recorded *x* and *y* coordinates of the movements were transformed into positional (*p*_*x*_, *p*_*y*_,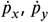), directional (*d*, ϑ, 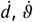) and non-directional (*s*) kinematics. Segments of EEG data with a length *w* prior to a time point *t*_0_ were extracted and labeled with the kinematics from *t*_0_ − 0.3 s. (b) The segmented EEG data was used to train and test a CNN designed according to the EEGNet architecture (***Lawhern et al., 2018***) which was modified to decode continuous values. The model consisted of a temporal convolution followed by a depthwise convolution and a separable convolution and was trained to decode hand kinematics from neural data during writing phases. Decoded kinematics were then cut to the length of the original trials from movement onset until 3.5 s after the movement onset. The time-series data of decoded kinematics affiliated with a written letter was used to train an SVM to classify the written letter from the decoded kinematics.

We extracted levels of statistical significance for results being better than chance for each participant by applying the procedure described by ***Müller-Putz et al. (2008***) based on the number of available trials per class. Apart from training models for the classification of all ten letters, we also trained models for the classification of subset *aenot* and subset *dfjsv*.

### Trajectory Decoding

The reconstruction of hand kinematics from EEG was achieved using the described EEGNet architecture adapted to the regression of multiple kinematics. We employed a mean squared error loss function and adapted the number of output neurons to the number of simultaneously decoded kinematics. Based on previous findings regarding the combination of directional and non-directional kinematics to improve the decoding of hand trajectories (***Mondini et al., 2020; Kobler et al., 2020c***), we opted to include directional (position) and non-directional (distance and speed) kinematics in the decoding. The *x* and *y* coordinates of the right index finger recorded with the motion capture system during hand movements were transformed into the following kinematics:

- Position-Based: Positions *p*_*x*_, *p*_*y*_ and velocities *v*_*x*_, *v*_*y*_ of the finger were directly obtained from the recorded coordinates and their derivatives.
- Distance-Based: Distance *d* and the derivative 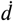 were obtained from the magnitude of the finger position. The angle ϑ was defined as the angle of the distance vector to the x-axis and 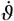 as the derivative of ϑ.
- Speed-Based: The speed *s* of the index finger was obtained from the magnitude of the velocity vector.

The relation of the kinematics with the coordinate frame and the letter trajectory is shown in Figure 2a. Since participants were instructed to start their movement from a comfortable position, the actual coordinates of the start were marginally different for every trial. To minimize variability within the trajectories, the coordinates were transformed to a starting point at a universal position [*x*_0_, *y*_0_] for each trajectory. Models were trained and tested on segments of EEG data filtered between 0.3–40 Hz and downsampled to 128 Hz (dataset 2). We extracted segments in windows of length *w* ∈ [0.4, 0.8, 1.2]s that included data prior to a time point *t*_0_ (i.e. [*t*_0_ − *w; t*_0_]) and were labeled with the corresponding value of the kinematic data at *t*_0_ − 0.3 s (see Figure 2a). This lag was introduced to incorporate information from reafferent neural signals related to the movement (***Seiss et al., 2002; Shibasaki and Hallett, 2006***) into the decoding of the movement kinematics. Windows were advanced with a stride of four samples starting from the movement onset until 3.5 s after the movement onset in each trial (i.e. *t*_0_ − 0.3 s ∈ [0.0; 3.5]s). This ensured that the complete movement during a trial was captured for most letters, since 99 % of all movements ended in this time interval. For all participants we used a two-times repeated three-fold cross-validation procedure which controlled for an equal ratio of letters per fold. In an inner loop, we further utilized a nested three-fold cross-validation to find the optimal window length. We selected the best window size based on the correlation of the decoded kinematics with the original kinematics of the validation test set. For each of the described kinematic sets, one model was trained to simultaneously predict the position-based, distance-based and speed-based kinematics. By storing and decoding the windowed data in order, we could resplit the decoded kinematics into the 3.5 s long trials corresponding to the letters. These trial-wise decoded kinematics were finally used to calculate the Pearson’s correlation coefficient *r* with the original kinematic and an average correlation was calculated for each of the nine kinematics. We extracted the statistical chance level for the correlation by testing the trained models on random phase surrogates of the test data. The approach used phase randomization by extracting the Fourier transformed data, randomizing the phase and applying the inverse Fourier transform to generate random data (***Lancaster et al., 2018***). This preserved certain statistical properties of the original signal such as the power spectrum while eliminating the original phase information. Since the surrogate test data thus did not contain any information related to the kinematics, we used the correlation between the actual kinematics and the kinematics predicted from surrogate data to estimate the chance level of the correlation. By calculating the correlation for 120 repetitions per kinematic, we estimated the chance level and the statistical level of 95 % confidence for correlation being significantly higher than chance.

### Two-Step Letter Classification

The classification of handwritten letters in the two-step approach was based on the previously decoded kinematics. For each round of the two-times repeated three-fold cross-validation, we utilized the model trained on the training set to decode the kinematics of every trial in the test and training set from EEG. We then trained a support vector machine (SVM) with the decoded kinematics of the training set and validated the classification accuracy on the decoded kinematics of the test set (see Figure 2b). We applied a sliding window approach similar to that used for direct classification by training SVM models on segments of kinematic data from each trial of lengths [*s* − *w; s*], *w* ∈ [0.5, 1.0, 1.5, 2.0, 2.5, 3.0, 3.5]s, *s* ∈ [*w*, 3.5]s with *s* being advanced with a stride of 0.1 s. Segments of the specified length and latency from all nine decoded kinematics were then used as features to the SVM. The model utilized a radial basis function kernel with a kernel coefficient selected according to 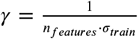 and squared l2 regularization with a penalty of 2.0. Kernel and regularization penalty were selected according to the optimal overall performance for participants at *w* = 3.5 s. For each subject, we obtained an individual classification accuracy by averaging over the results obtained from the cross-validation procedure. Classification accuracies for the subsets *aenot* and *dfjsv* were obtained by training and testing models on the subsets of decoded trajectories for the specific letters. We further investigated the importance of kinematics for the classification of letters by applying randomized permutation to the kinematic features (***Altmann et al., 2010***). For each kinematic, features were permuted across trials to break the connection between the data and label for the specific kinematic. The model was then evaluated on the permuted data and differences to the baseline from classification of non-permuted kinematics were calculated. By repeating this procedure multiple times for each kinematic, the importance of each feature for the classifier could be estimated. Higher differences between the accuracy obtained with the permuted set and the baseline demonstrated greater importance of the feature. Since the decoded kinematics were highly correlated as they included derivatives of the same kinematic, we employed a clustering method and calculated the feature importance for each kinematic inside the cluster. Clusters were selected to contain either the kinematic or its derivative, i.e. *d* and ϑ would not be combined with 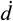 or 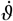. We thus calculated the importance for four groups (*d*, ϑ, *s, p*_*x*_, *p*_*y*_ /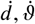, *s, p*_*x*_, *p*_*y*_/ *d*, ϑ, *s, v*_*x*_, *v*_*y*_ and 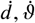, *s, p*_*x*_, *p*_*y*_). The final feature importance was then calculated from the average of the importances per kinematic. We calculated feature importances per participant for the set of all letters as well as the subsets *aenot* and *dfjsv*.

## Results

### Neural Correlates

Results for the analysis of low-frequency neural correlates of handwritten letters are shown in ***Figure 3***. MRCPs were visible for all letters and showed a centralized contra-lateral distribution in channels affiliated with the motor cortex (***Figure 3***a). The grand-average MRCPs for individual letters exhibited differences in the amplitude and latency as shown for the letters *o* and *v* in ***Figure 3***b. There was no significant correlation between the minimum amplitude of the MRCP around the movement onset and the average speed or duration of the written letters at electrodes C3, C1, Cz, C2 and C4. Similarly, we found no significant correlation between the average speed or duration and the rebound rate of the MRCPs. The results from the analysis of individual differences in low-frequency time-domain data between letters revealed significant variations at different time stages (***Figure 3***c). Around the movement onset (−0.5 s–0.5 s, lower left triangle), significant differences could be observed in central channels and in occipital channels contralateral to the movement. Letters *a, d, o* and *s* showed a higher number of channels with significant differences compared to other letters. 0.5 s–1.5 s after the movement onset (upper right triangle), fewer significant differences were visible in the neural data between letters. While these differences occurred mostly in central channels, no clear lateralization and no significant differences were found in occipital regions at this time stage.

**Figure 3.**
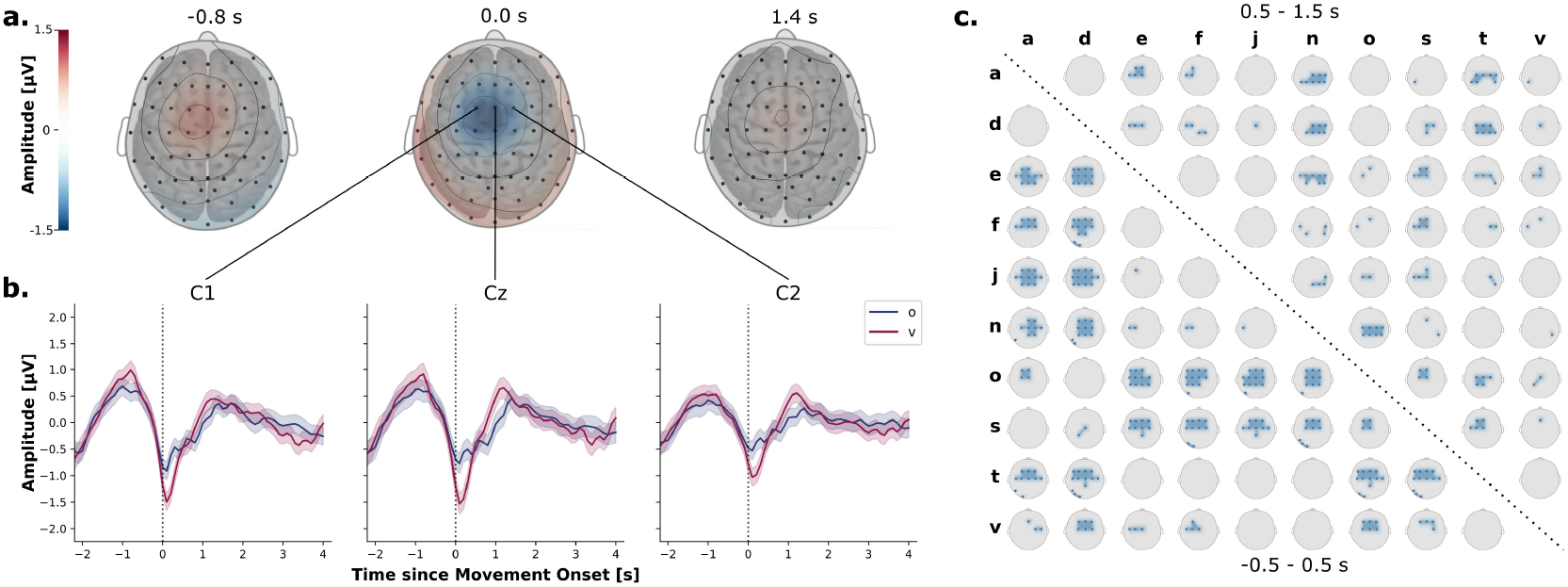
(a) Topographical plots of grand-average MRCPs from low-frequency EEG in sensor-space 0.8 s before, during and 1.4 s after the movement onset. The topographical plots show the amplitude distribution on the scalp with the brain for orientation. (b) Grand-average MRCPs with 95 % confidence interval for letters *o* and *v* at channels C1, Cz and C2. The movement onset is indicated by a dotted line at 0.0 s. (c) Statistically significant channels for the pairwise comparison of low-frequency EEG data around and after the movement onset for the different letters. The lower left triangle shows the significant channels for the time period of −0.5–0.5 s and the upper right triangle shows the significant channels for the time period of 0.5–1.5 s.

### Letter Classification

The sliding-window approach revealed an optimal window length of *w* = 2.0 s for sLDA and EEGNet models. The overall curves of the sliding-window approach composed from the average classification results of all subjects obtained at different latencies *l* to the movement onset in ***Figure 4***b depict the ideal latency for classification for each model. The highest classification accuracy for all ten classes was obtained with a latency of 2.0 s for the sLDA and 1.7 s for the EEGNet with an accuracy of (mean ± standard deviation) 23.1 ± 6.8 % and 22.2 ± 8.6 %, respectively. For subsets *aenot* and *dfjsv*, similar analyses revealed highest accuracies of 35.3 ± 9.0 % (latency 2.0 s, *aenot*) and 39.0 ± 9.6 % (latency 2.0 s, *dfjsv*) for the sLDA and 33.5 ± 9.0 % (latency 1.9 s, *aenot*) and 38.3 ± 11.4 % (latency 2.1 s, *dfjsv*) for the EEGNet. These results are summarized in Table 1. No significant differences between the models were found for each of the three sets as assessed with a paired t-test.

**Table 1.**
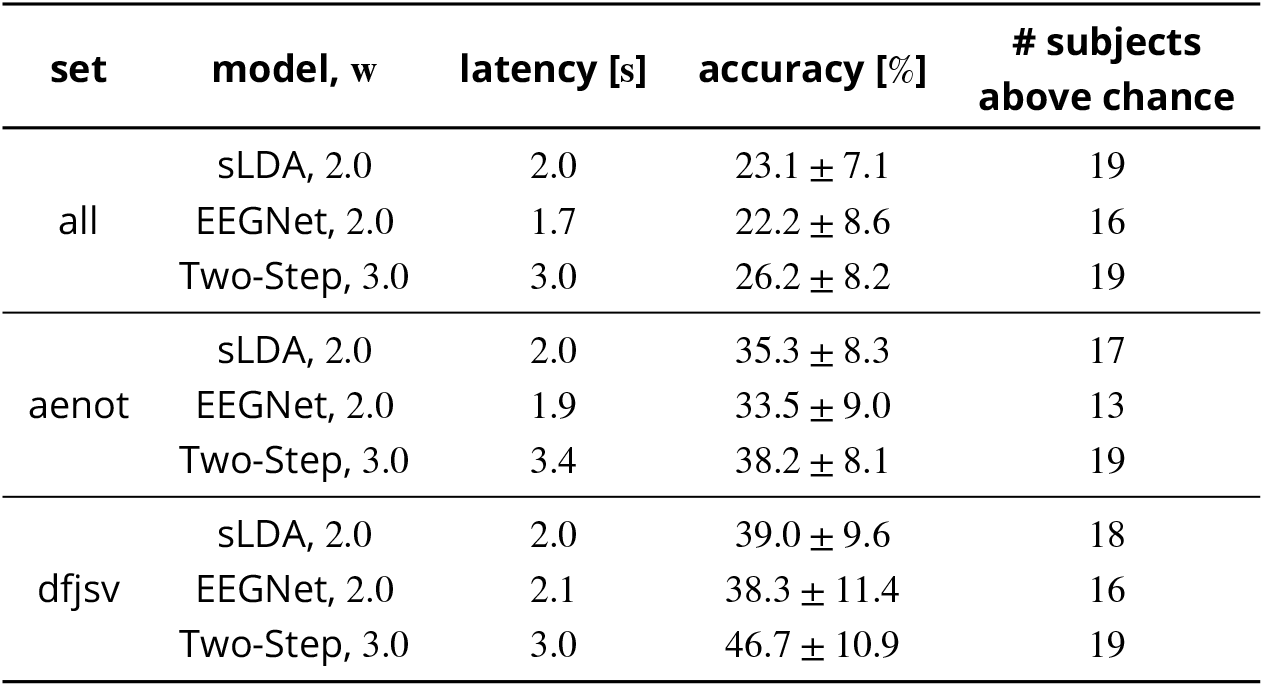
Overall accuracies for the different models and letter sets for the best performing window length and latency. The last column shows the number of subjects with results significantly above chance.

| set | model, w | latency [s] | accuracy [%] | # subjects above chance |
| --- | --- | --- | --- | --- |
| all | sLDA, 2.0 | 2.0 | 23.1 ± 7.1 | 19 |
|  | EEGNet, 2.0 | 1.7 | 22.2 ± 8.6 | 16 |
|  | Two-Step, 3.0 | 3.0 | 26.2 ± 8.2 | 19 |
| aenot | sLDA, 2.0 | 2.0 | 35.3 ± 8.3 | 17 |
|  | EEGNet, 2.0 | 1.9 | 33.5 ± 9.0 | 13 |
|  | Two-Step, 3.0 | 3.4 | 38.2 ± 8.1 | 19 |
| dfjsv | sLDA, 2.0 | 2.0 | 39.0 ± 9.6 | 18 |
|  | EEGNet, 2.0 | 2.1 | 38.3 ± 11.4 | 16 |
|  | Two-Step, 3.0 | 3.0 | 46.7 ± 10.9 | 19 |

**Figure 4.**
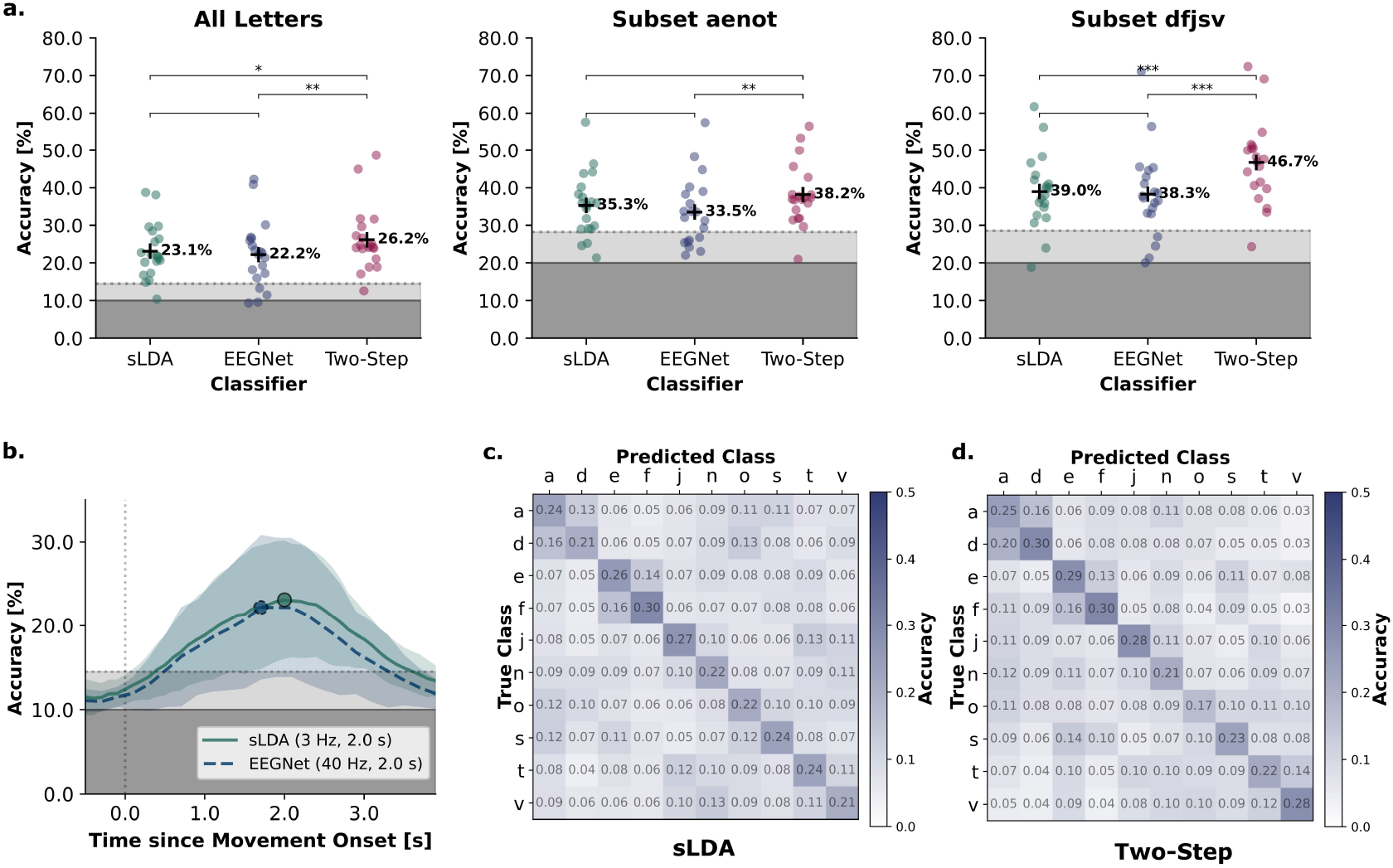
(a) Final results obtained for the classification accuracies of letters using the proposed models and different letter subsets. The statistical significance from paired t-tests after Bonferroni correction between the classification results is indicated by asterisks (*: *p* ≤ 0.05, **: *p* ≤ 0.01, ***: *p* ≤ 0.001). The dark grey area represents the theoretical chance level, the light grey area the level of statistical significance for results being better than chance. (b) Comparison of the resulting curve of windowed letter classification using sLDA and EEGNet models for a window length of 2.0 s and different latencies. The optimal latencies to achieve the highest classification performances are marked with a circle. (c) Overall confusion matrix for the classification of all letters using the best performing model of the direct classification (sLDA with a window length of 2.0 s and a latency of 2.0 s). Values are normalized along the rows (true classes). (d) Overall confusion matrix for the classification of all letters using the two-step approach and a window length of 3.0 s at a latency of 3.0 s for the SVM. Values are normalized along the rows (true classes).

All models performed significantly better for the subset *dfjsv* than for the subset *aenot* (*p*_*sLDA*_ = .017, *p*_*EEGNet*_ = .009). The statistical chance level indicated by a light grey area in ***Figure 4***a and b, compared to the theoretical chance level of 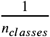 indicated by a dark grey area, amounted to 14.5 % for the set of all letters, 28.3 % for the subset *aenot* and 28.6 % for the subset *dfjsv*. Out of 20 participants, one participant did not perform higher than chance for the sLDA on the set of all all letters and three/two participants for the subsets *aenot*/*dfjsv*, respectively. With the EEGNet model, four participants did not perform higher than chance on the set of all letters, seven participants on subset *aenot* and four participants on subset *dfjsv*. The two best performing subjects achieved accuracies of 38.0 %/38.7 % and 42.2 %/40.8 % for all letters, 57.6 %/46.3 % and 57.5 %/48.2 % for the subset *aenot* and 57.6 %/46.3 % and 56.4 %/71.1 % for the subset *dfjsv* with sLDA and EEGNet. A confusion matrix summarizing the performance of the sLDA classifier for all subjects for the complete letter set is provided in ***Figure 4***c. The matrix is normalized along the rows (true classes) and shows the results of the predicted classes utilizing the best performing sLDA model with a window length of 2.0 s at a latency of 2.0 s. Confusion matrices for all letter subsets and the EEGNet classifier can be found in ***Figure S1*** together with the results for other window lengths obtained in the sliding-window approach in ***Figure S2***.

### Trajectory Decoding

The nested cross-validation procedure selected the optimal window length to be 0.4 s in 12.5 %, 0.8 s in 43.3 % and 1.2 s in 44.2 % of the cases. ***Figure 5***a shows the recorded and reconstructed distance (red), speed (green) and y-position (blue) trajectories of three exemplary letters (*v, a, o*). The reconstructed kinematics of 3.5 s length were correlated with the original kinematics obtained from the recorded movements. The average correlations of all trials per participant and kinematic are shown in ***Figure 5***c. The overall average correlations per kinematic, as indicated by a black cross, were (mean ± standard deviation) 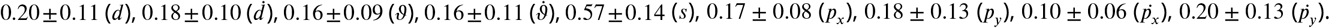. Statistical chance levels, indicated by a light gray area, ranged between 0.038 and 0.046 for different kinematics. The reconstructed speed achieved a significantly (*p* < 0.001 after Bonferroni correction for multiple comparisons) higher correlation with the original kinematic than was achieved for any of the other kinematics as assessed with a paired t-test. Similarly, the mean squared error between z-scored kinematics was significantly lower for speed compared to all other kinematics (see ***Figure S3***). Dependent on the kinematic, between zero (speed) and four (velocity along the x-axis) participants did not achieve a correlation significantly greater than chance.

**Figure 5.**
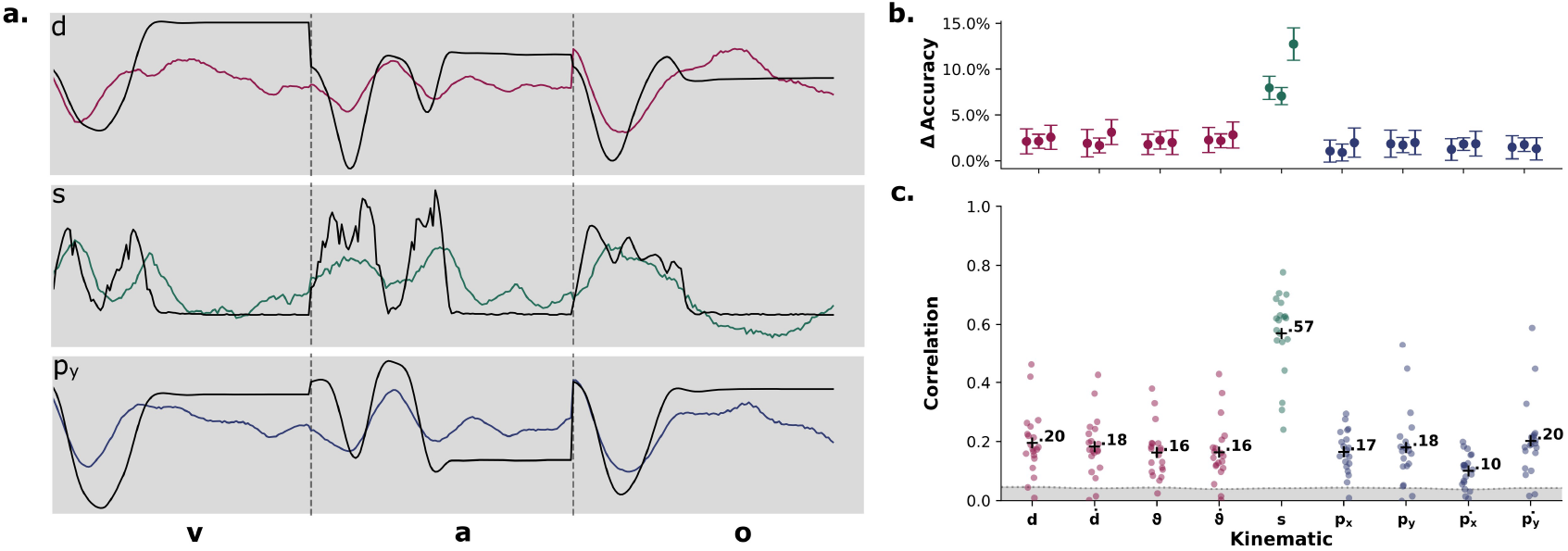
(a) Exemplary recorded (black) and decoded (colored) kinematics of single trials for letters *v, a* and *o* of the kinematics distance (*d*), speed (*s*) and position along the y-axis (*p*_*y*_). The correlations for *d, s* and *p*_*y*_ of the specific subject were 0.46, 0.70 and 0.53, respectively. (b) Importance of the kinematics for the classification performance of the SVM for different sets of letters as calculated from a permutation approach (for each kinematic from left to right: all letters, subset *aenot*, subset *dfjsv*). The standard deviation is shown and calculated from feature importances achieved for single participants. (c) Average correlation from all trials for the kinematics per participant (individual dots) and overall (black cross). The light grey area represents the chance level of the correlation for the different kinematics. Kinematics are displayed in color based on being derived from distance (red), speed (green) or position (blue).

### Two-Step Letter Classification

The classification of letters from decoded kinematics using an SVM resulted in an overall accuracy of (mean ± standard deviation) 26.2 ± 8.2 % for the set of all letters, 38.2 ± 8.1 % for the subset *aenot* and 46.7 ± 10.9 % for the subset *dfjsv* at an optimal window length of 3.0 s with latencies of 3.0, 3.4 and 3.0 s, respectively. The model performed significantly (*p* < .001) better for the subset *dfjsv* than for the subset *aenot*. Compared to the classification results obtained with sLDA and EEGNet from direct classification, the two-step approach resulted in a performance increase of 3.1 % (all letters), 2.9 % (subset *aenot*) and 7.7 % (subset *dfjsv*). Classification accuracies were significantly higher than the results achieved with either sLDA or EEGNet for all letters (*p*_*sLDA*_ = .019, *p*_*EEGNet*_ = .001) and for the subset *dfjsv* (*p*_*sLDA*_ < .001, *p*_*EEGNet*_ < .001) and significantly higher than those achieved with the EEGNet for the subset *aenot* (*p*_*sLDA*_ = .136, *p*_*EEGNet*_ = .006) as assessed with paired t-tests. *p*-values were corrected for multiple comparisons using Bonferroni correction. ***Figure 4***a shows a comparison between the results of all models. Out of 20 participants, one subject did not achieve a classification accuracy higher than chance for all three sets of letters. The two subjects achieving the best performances (similar subjects as above) obtained accuracies of 44.9 %/48.6 % (all letters), 56.5 %/53.2 % (subset *aenot*) and 69.1 %/72.4 % (subset *dfjsv*). ***Figure 4***d shows the overall confusion matrix for the set of all letters utilizing the two-step classification approach. The matrix is normalized along the rows (true classes). The importance of kinematics to the letter classification investigated with a permutation approach is given in ***Figure 5***b as the difference between the accuracy achieved with permuted features and the non-permutated baseline. The feature importance is shown for the three different sets of letters and attributes the highest importance to the speed accounting for (mean ± standard deviation) 8.0 ± 1.0 % (all letters), 7.1 ± 1.3 % (subset *aenot*) and 12.7 ± 1.8 % (subset *dfjsv*) of the performance. Colors indicate which kinematics are derived based on distance (red), speed (green) or position (blue). Confusion matrices for all subsets can be found in ***Figure S1*** and results for all window lengths are shown in ***Figure S2***.

## Discussion

In the current work we showed for the fist time the application of continuous movement decoding from non-invasive EEG for the classification of handwritten letters. We also demonstrated how a two-step approach of decoding and subsequent classification can increase classification performances for letters with trajectories chosen based on high discriminability. Results suggest that the classification of handwritten letters from EEG is generally feasible and provide an overview of the decoding of movement kinematics of short, complex hand movements, revealing significantly higher correlations for the movement speed than for other kinematics.

Although the overall classification accuracies are modest and not in a range to enable suitable communication for patients, single participants reached high performances in the classification of a subset of five letters (*dfjsv*). These performances of up to 72 % accuracy approach the requirement of 75 % accuracy in communication devices for patients (***Pfurtscheller et al., 2005***). Compared to results of 76.8–97.0 % obtained by ***Pei and Ouyang (2021***) for the classification of nine characters, the performances achieved in the current study are substantially lower. However, ***Pei and Ouyang (2021***) did not correct for eye artifacts during the classification process and did not adapt the study design to minimize eye movements. This could have artificially improved the classification accuracy since EOG signals have been shown to influence performances, e.g., in the decoding of three-dimensional trajectories (***Kim et al., 2015***). Center-out tasks employing EEG for the classification of hand movements in four opposite directions have been shown to achieve performances in the range of on average 55.9–80.2 % (***Kobler et al., 2020a; Shiman et al., 2017; Robinson et al., 2013***). In a decoding approach closely related to the current study, ***Úbeda et al. (2017***) attempted the classification of center-out movements based on continuous decoding of hand motions from EEG, reaching average accuracies of 51.3 % or 52.3 % for four and 29.0 % for eight directions. Although these performances surpass those documented within our study, they confirm that our results fit within the normal range of performances for EEG-based classification.

While the sLDA model performed better than the EEGNet for all three sets, differences were not significant. The EEGNet did perform equally well or better than the sLDA for the two highest performing subjects, indicating the similar capability of both models for the classification of handwritten letters from EEG. It is of interest to notice the difference in features utilized by the models: while the input to the sLDA consisted of low-frequency EEG, the EEGNet employed data from a broad frequency spectrum and learned appropriate frequency-specific features from the training data. Since these models performed nearly equally well, this could suggest that no additional information regarding the written letter was utilized from higher frequency ranges. This is in accordance with a large number of studies solely using low-frequency components for the direct classification of movements from EEG including hand (***Bressan et al., 2021; Sburlea et al., 2021; Úbeda et al., 2017; Ofner et al., 2017a; Robinson et al., 2013***) and finger (***Kobler et al., 2020c; Mondini et al., 2020***) movements. ***Borra et al. (2023***) similarly found low-frequency components to be most informative for the decoding of movement kinematics. When comparing the confusion matrices for sLDA and EEGNet (see ***Figure S1***) with the significant differences in low-frequency EEG at ±0.5 s around the movement onset between letters, high similarity between patterns of few significant differences and decreased decodability can be observed. Letters without any significant difference in low-frequency EEG like *a*/*d* or *e*/*f* achieved worse decoding performances than letters with more significant differences like *a*/*d* with *e*/*f* /*j*/*n*. This connection can be observed both in EEG-Net and sLDA confusion matrices, suggesting a connection of both models to the low-frequency components of EEG.

Further, investigation of low-frequency time-domain data around the movement onset showed clearly distinguished MRCPs localized contralateral to the movement as has been described extensively throughout literature (***Shakeel et al., 2015***). In accordance with ***Gu et al. (2009***) we did not find any significant correlation between the writing speed or duration and the amplitude of the negative deflection. However, we also did not find any significant correlation between the average speed of the written letters and the rebound rate, which contrasts their finding of increased rebound rates correlating with higher speeds of movements. These results might be explained by the higher complexity of the movements executed in the current study and the changes in writing speed within each letter. However, also the comparatively low numbers of approximately 46 trials per subject and letter utilized for the generation of MRCPs in our study may have contributed to the outcome. The statistical evaluation of differences in neural data between letters found the highest number of significant differences in channels contralateral to the movement in the temporal window of ±0.5 s around the movement onset. This indicates an involvement of motor areas and suggests the movement as a confounding factor for differences in the neural signals compared to visual or cognitive processes. As the experiment was explicitly designed in a way that no visual input was shown to the participants during the execution of the movement, these findings approve the design of the study to be mostly independent from visual input. Additionally, frontal channels were not found to be significantly different, confirming the independence of the neural signals from eye movements connected to the written letter. The lateralization of neural differences is in accordance with literature: ***Schwarz et al. (2020***) found significant differences between grasp types within 0.5 s after the movement onset over contralateral central areas and similar results have been found by other studies (***Ofner et al., 2017a; Iturrate et al., 2018; Agashe et al., 2015***). The significance of occipital channels in some comparisons can be explained by the projection of MRCPs to the lateral channels through the re-referencing to the common average. When calculating significant differences for low-frequency time-domain signals without any re-referencing, such occipital differences do not occur. Unlike ***Mondini et al. (2024***) who found activity related to the movement in parieto-occipital areas, the current study did not find significant differences in these areas. This might be related to the different study design, since ***Mondini et al. (2024***) employed a pursuit tracking task which utilized hand-eye coordination while our study was explicitly designed to exclude visual input. For the time segment between 0.5–1.5 s after the movement onset which includes the termination of the writing movement for the first 30 % of motions, significant differences occurred similarly in central regions. However no clear lateralization was evident. These significant differences in time-series neural data could be related to the continuous movement executed during this time since they have not been found to be significant for single movements in other studies (***Schwarz et al., 2020***). This is also suggested by the optimal window length for direct classification of letters from EEG including data until around 2.0 s after the movement onset.

Optimal window length and latency obtained from the sliding-window approach were comparable for both sLDA and EEGNet in all sets, incorporating data from the movement onset until 1.7– 2.1 s after the onset. Compared to a study on the classification of single movements which found optimal window lengths of 1.4 s (***Ofner et al., 2019***), this window length is considerably longer. Interestingly, the optimal window was still substantially shorter than a large number of the executed movements with 61 % of all movements being completed during this time. In contrast to the direct classification, the sliding-window approach used for the classification of letters from decoded kinematics revealed longer optimal windows of 3.0 s which incorporated the complete movement in 95 % of the trials.

The improvement in accuracy for all sets of letters obtained with the two-step approach suggests that the classification of handwritten letters can be enhanced through the decoding of continuous kinematics. While all models showed an increased accuracy in subset *dfjsv* compared to subset *aenot*, the increase was substantially higher for the two-step approach. This suggests that the selection of letters based on higher discriminability between the trajectories could be exploited more by the concatenation of kinematic decoding with subsequent classification than by direct classification from EEG. The analysis of feature importances and the achieved correlation of the reconstructed and original kinematics showed significantly higher correlation for speed than for any other kinematic as well as a substantially higher dependence of the SVM on speed during classification. Although the subset *dfjsv* was not originally selected based on differences in writing speed, a comparison of correlations of speed trajectories between letters (see ***Figure S4***) showed an overall higher correlation between speed trajectories for subset *aenot* (average correlation of 0.76) than for subset *dfjsv* (average correlation of 0.49). This suggests that the differences in speed trajectories between letters in subset *dfjsv* compared to *aenot* could be responsible for the increase in classification performance. As indicated also by the not significant influence of speed on the MRCPs, the patterns between significant differences in low-frequency EEG and the correlation of speed trajectories between letters did not show high similarities. Furthermore, the fact that the direct classification of letters from EEG using the EEGNet did not achieve a similarly high accuracy as that using the two-step approach, even though both models utilized similarly processed EEG data (dataset 2), indicates that direct classification was unable to utilize information encoded in the tuning of neural data to speed when only provided with the label of the written letter. However, it seems that the utilization of this information is essential for the improvement of classification accuracies. A selection of letters based on differences in speed of the trajectories could thus further improve the performance of the two-step approach while it might not benefit the direct classification results.

A similar tuning of neural activity to the writing speed of the movement during handwriting was reported by ***Willett et al. (2021***) in their study on the classification of handwritten letters from invasive recordings with microelectrode arrays. This is in contrast to studies on the non-invasive decoding of movement kinematics, which reported higher correlations between decoded and original kinematics for directional (position and velocity in *x* and *y* direction) than non-directional (distance and speed) kinematics (***Kobler et al., 2020c***). However, the type of movement could be relevant to the higher correlation found in this study, since ***Kobler et al. (2020c***) investigated continuous movements with a duration of 16 s during a pursuit tracking task while the current study investigated movements with an average duration of 1.9 ± 0.61 s. In general, most previous studies on non-invasive decoding of hand movements employed pursuit-tracking tasks or related paradigms such as center-out tasks in which the participants had to draw simple, straight lines. This adaptation of simple or externally dictated movements does not allow for any planning of complex motions, thereby possibly altering the involved neural dynamics. A study investigating the effect of *goal* vs. *no-goal* directed movement found significant differences in low-frequency EEG for these two conditions (***Pereira et al., 2017***). Thus, it is reasonable to assume that the goal-directedness introduced by the self-governed writing of the given letter could play a role in the decoding of kinematics and may be responsible for the differences in achieved correlation. The substantially lower correlation obtained for directional kinematics compared to (***Kobler et al., 2020c***) might however also be explained by changes in the neural data due to differences in the paradigm: while ***Kobler et al. (2020c***) investigated hand movements, the current study investigated movements of an individual finger.

Regarding communication speed and accuracy, the proposed method is unable to reach the performances of classifiers utilizing neural signals recorded with invasive microelectrode arrays (***Willett et al., 2021***). However, since such recordings require lengthy preparation phases and patients undergoing surgery, EEG can offer an alternative to people unwilling to expose themselves to risks of surgery or during the waiting phase for an implantation. Since the proposed method relies on similar neural underpinnings as methods reaching high performances with intracranially recorded signals, it could also function as an early training or screening process for patients prior to implantation.

The main limitations of the current study concern the offline analysis of the recorded data and the unclear transferability of results to locked-in patients. While most preprocessing steps, including the eye artifact correction, can also be used in an online adaptation of the paradigm, the influence of residual artifacts when not correcting these with the usage of ICA would need to be considered in an online paradigm. However, inspected components suggested that only a limited influence of eye or movement artifacts remained before applying ICA, thus indicating a minimal influence on the decoding performance in an online approach. The adaptation of the paradigm to locked-in patients poses considerably greater problems, since the presented paradigm relies on executed movements. While studies have reported the successful decoding of attempted movements from spinal cord injured participants (***Pulferer et al., 2022; Ofner et al., 2019; Lopez-Larraz et al., 2012***), it is currently unclear whether performances suited to the classification of handwritten letters could be obtained from attempted movement with the proposed two-step approach. Further, the generation of training data for the decoding of kinematics in the two-step approach remains problematic for locked-in patients. Future work will address these limitations by adapting the proposed paradigm.

## Conclusion

The current study demonstrated for the first time the utilization of continuous decoding of hand kinematics for the classification of handwritten letters from EEG. While this two-step approach yielded significant but moderate improvements in performance for a set of ten letters compared to direct classification using sLDA and EEGNet models, the increase in accuracy for a subset of five letters with highly discriminable trajectories was substantial. We assume a tuning of neural data to the writing speed based on the significantly higher correlation of reconstructed with original kinematic for speed than for other kinematics as well as the high feature importance assigned to the speed kinematic during letter classification. These findings are in accordance with previous studies utilizing invasive recordings and show that the general concepts can be transferred also to non-invasive signals. The results suggest that the classification accuracy could be further increased by selecting most discriminable letters based on differences in the speed of the writing trajectory. Importantly, this study showed that the proposed two-step approach possesses a greater capacity for the utilization of optimized writing trajectories than models classifying letters directly from EEG, indicating the benefit of the proposed approach for the improvement of classification for handwritten letters. Further work will focus on the utilization of speed differences for the improvement of classification accuracies to enable performances suitable for actual communication. While some participants already showed performances in the required range, this was not the case for the majority of subjects. Finally, this study showed that the classification of handwritten letters from non-invasive EEG is generally feasible and could be used as a way of communication for patients in a locked-in state in the future. The spelling of complete words from decoded letters could then also be aided by the introduction of a language model to increase the overall performance of the model.

## Acknowledgments

We thank all participants for their contribution to this study and acknowledge Kyriaki Kostoglou, Johanna Egger and Patrick Suwandjieff for their valuable support and discussions. This work was funded by the European Union’s HORIZON-EIC-2021-PATHFINDER CHALLENGES program under grant agreement No 101070939 and by the Swiss State Secretariat for Education, Research and Innovation (SERI) under contract number 22.00198.

---

**Supplemental Figure S1.**
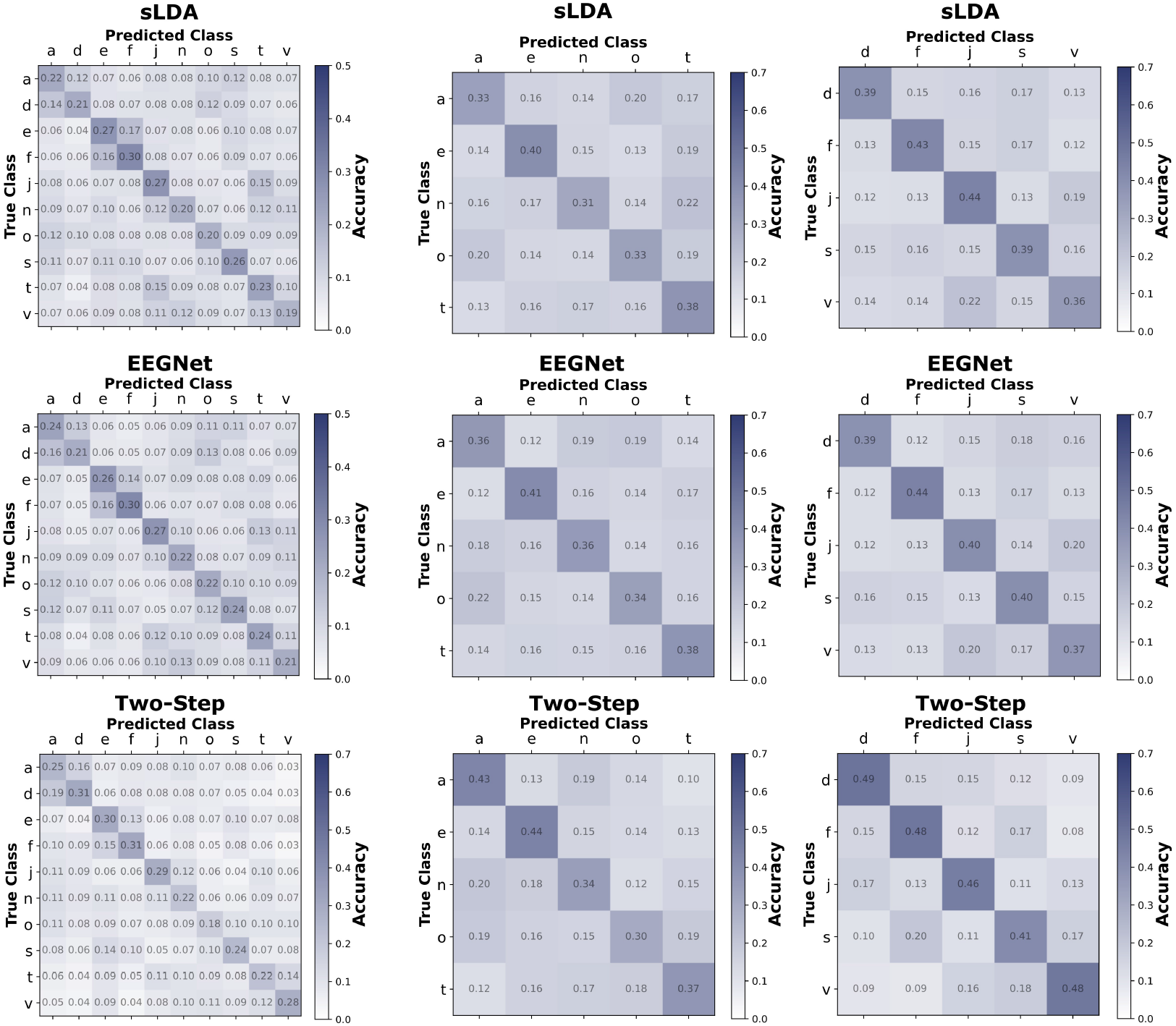
Confusion matrices for the classification of all letters, subset *aenot* and subset *dfjsv* for models of direct classification (sLDA and EEGNet) and the two-step approach. The confusion matrices show results for the best performing window length and latency. Confusion matrices are normalized along the rows (true classes).

**Supplemental Figure S2.**
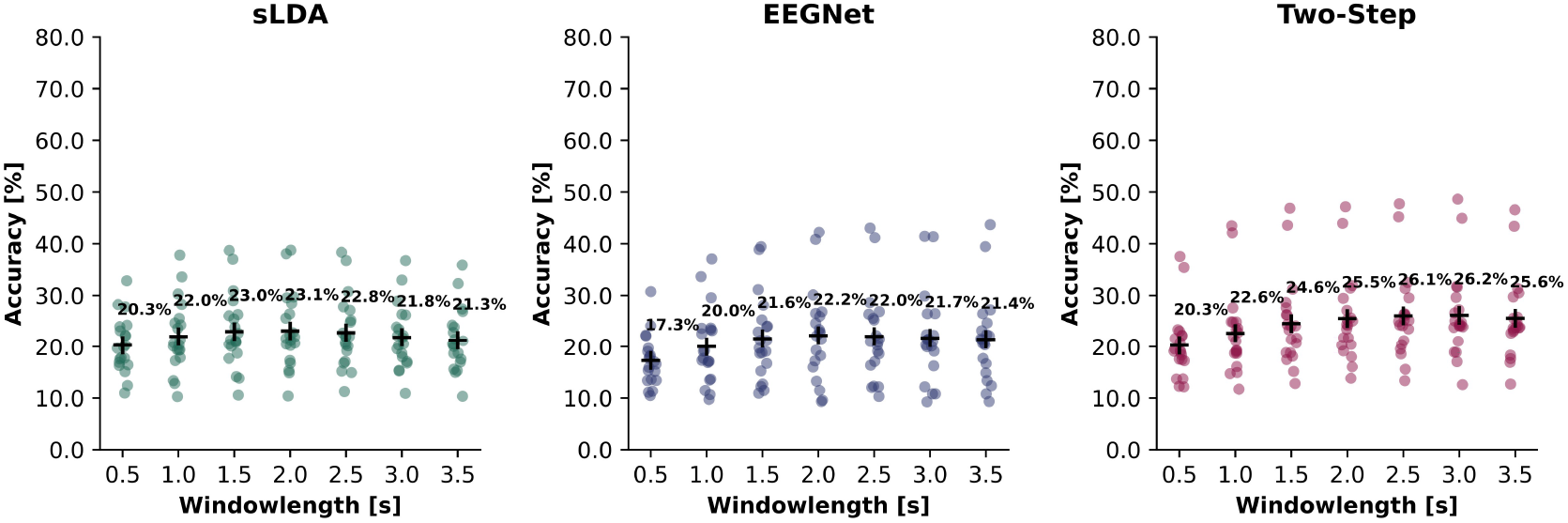
Results of the sliding-window approach for different window lengths at optimal latencies. Results are displayed for sLDA and EEGNet for direct decoding from EEG and for the two-step approach from kinematics decoded from neural data. The optimal latencies for which results are depicted were found to be [1.0, 1.3, 1.8, 2.0, 2.4, 2.9, 3.0]*s* for sLDA, [0.9, 1.3, 1.7, 1.7, 2.3, 2.3, 2.7]*s* for EEGNet and [1.4, 1.7, 2.1, 2.5, 2.6, 3.0, 3.5]*s* for the two-step approach.

**Supplemental Figure S3.**
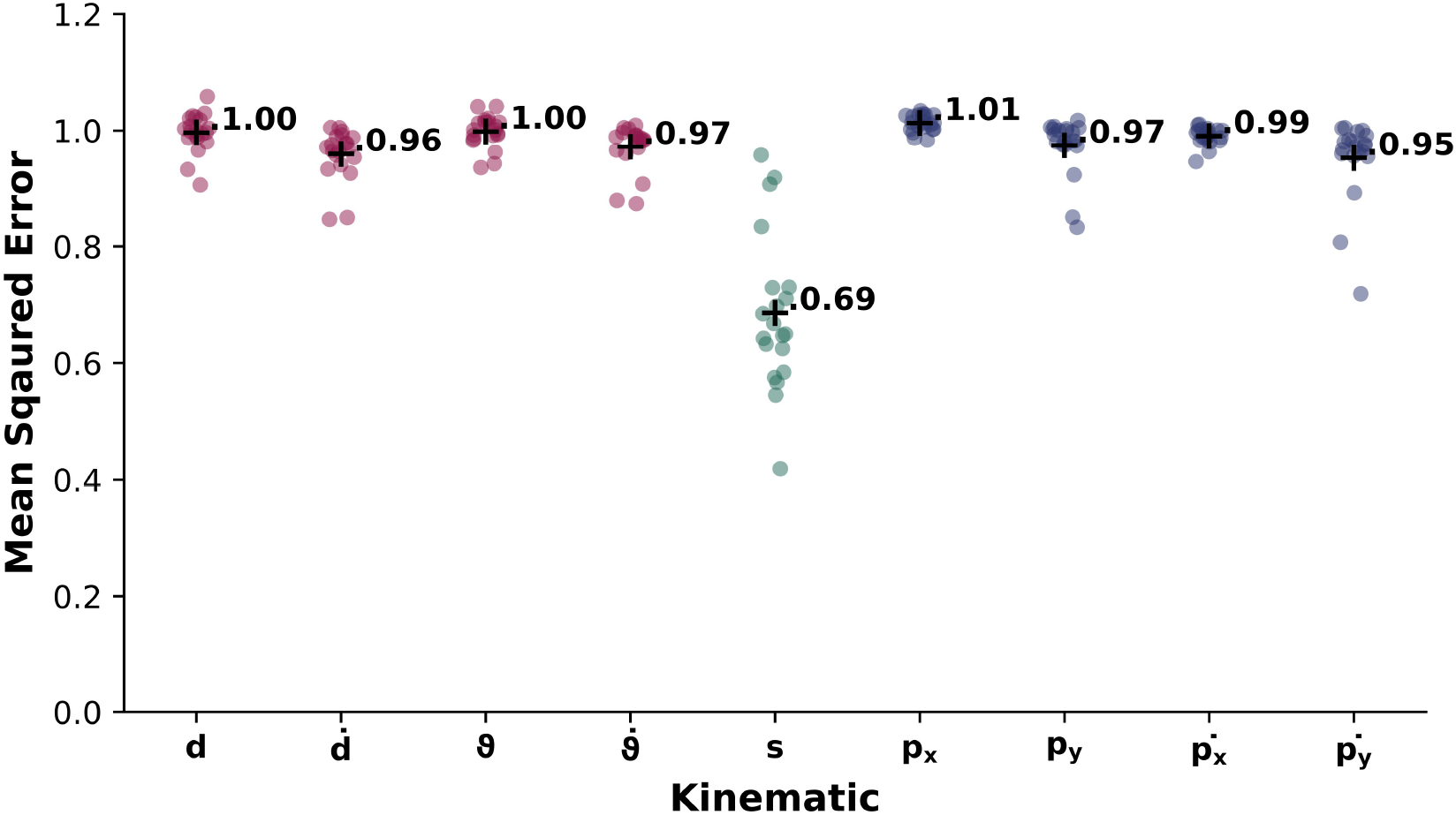
Mean squared error for all participants and kinematics calculated between the decoded and original kinematics of each trial. Kinematics were z-scored with the mean and standard deviation of the original kinematics to enable comparisons between kinematics. The black cross indicates the overall mean value.

**Supplemental Figure S4.**
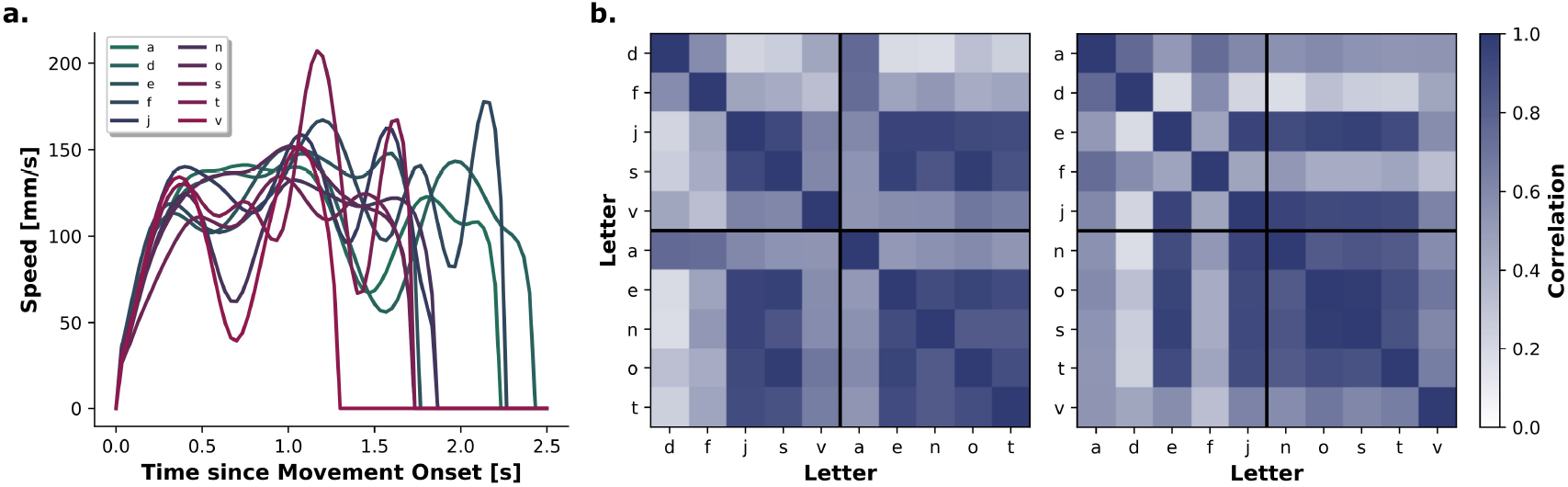
(a) Average speed trajectory per letter from all trials after interpolation to the average length. (b) Correlation of speed between letters from interpolated trajectories. The left matrix is ordered according to the subsets of letters, the right matrix shows letters in alphabetical order. In the left correlation matrix, the upper left square corresponds to the correlation of the speed within the subset *dfjsv*, the lower right square to that of subset *aenot*. The average correlation within the upper left square (without trace) is 0.49, the average correlation within the lower right square is 0.76.

